# DNA methylation profile is not inherited in offspring of a short-lived annual fish

**DOI:** 10.64898/2026.07.28.741221

**Authors:** Malahat Dianat, Lisandrina Mari, Dagmar Čížková, Milan Vrtílek

## Abstract

Parental ageing can influence offspring through non-genetic mechanisms. The contribution of epigenetic inheritance parental effects still remains poorly understood. DNA methylation is a widespread regulator of gene expression that changes during development and ageing and can also act as a mediator of intergenerational effects. We tested whether age-related changes in parental DNA methylation are transmitted to offspring in the short-lived turquoise killifish (*Nothobranchius furzeri*, Cyprinodontiformes). Using reduced-representation sequencing, we quantified genome-wide DNA methylation and examined methylation dynamics at individual loci. The overall proportion of methylated CpG sites increased during early ageing but declined at later ages, revealing a non-linear trajectory with substantial among-individual variation. Despite these age-related changes, we found no evidence that parental methylation patterns were transmitted to offspring, either at the genome-wide level or at individual loci. Our findings indicate that although DNA methylation undergoes pronounced age-dependent remodelling in adult killifish, these changes are not detectably inherited by the next generation. We discuss these results in the context of epigenetic inheritance and ageing in vertebrate model systems.

## Introduction

Senescence is an individual and complex process characterized by progressive deterioration of physiological functions and fitness with advancing age (Levine, 2013; López-Otín *et al*., 2013). Within a population, individuals of the same age usually vary in their physiological state, i.e. their biological age (Jackson *et al*., 2003). One of the central components of senescence and biological ageing is the shift in epigenetic regulation of cellular processes (López-Otín *et al*., 2013; Pal & Tyler, 2016; Horvath & Raj, 2018). Beyond their role in regulating gene expression during ageing, epigenetic marks also constitute a layer of inheritance (Fitz-James & Cavalli, 2022).

The epigenetics of ageing therefore represents a promising avenue for better understanding of the sources of individual variation in senescence and its potential mitigation. DNA methylation, the addition of methyl group to cytosine residues, is an evolutionarily conserved and ontogenetically dynamic cellular mechanism to regulate gene expression (Zemach *et al*., 2010; Klughammer *et al*., 2023; Li *et al*., 2024). Both passive (epigenetic erosion) and active changes in methylome occur during ageing (Le Clercq *et al*., 2023). Systematic alterations in DNA methylation associated with ageing provide basis for epigenetic clocks (Lu *et al*., 2023). These clocks can then be used to precisely estimate not only actual chronological age (Horvath & Raj, 2018; Le Clercq *et al*., 2023; Arai & Inoue-Murayama, 2026) but also to measure age-related physiological decline (senescence) and approaching death (Pal & Tyler, 2016; Levine *et al*., 2018; Giannuzzi *et al*., 2024). Some studies indicate that early-life dynamics can modulate epigenetic ageing trajectories, highlighting both conserved mechanisms and context-dependent variation in epigenetic ageing (Parrott & Bertucci, 2019; Bertucci *et al*., 2021).

There is a mixed evidence for potential inter-generational transmission of epigenetic ageing across vertebrates (Heard & Martienssen, 2014; Skvortsova *et al*., 2018; Fitz-James & Cavalli, 2022). In human and mice, global demethylation of parental genomes during embryogenesis prevents large-scale accumulation of epigenetic alterations across generations but some regions may still escape the erasure (Hackett *et al*., 2013; Skvortsova *et al*., 2018; Kremsky & Corces, 2020). In zebrafish (*Danio rerio*), early embryonic DNA methylation reprogramming is incomplete and offspring DNA methylation pattern is skewed towards paternal methylome (Jiang *et al*., 2013; Potok *et al*., 2013; Skvortsova *et al*., 2019). In another model laboratory fish, the medaka (*Oryzias latipes*), DNA methylation inheritance is closer to the mammalian type – largely erased during embryogenesis (Wang & Bhandari, 2019), and the same is true for embryos mangrove rivulus (*Kryptolebias marmoratus*) (Fellous *et al*., 2018). In general, epigenetic inheritance is considered a variable and context-dependent process (Heard & Martienssen, 2014; Perez & Lehner, 2019; Heckwolf *et al*., 2020; Fitz-James & Cavalli, 2022). To address the potential epigenetic inheritance an experimentally tractable vertebrate model is required.

Turquoise killifish (*Nothobranchius furzeri* Jubb 1971, Cyprinodontiformes) is an emerging model for studying ageing due to its exceptionally short natural lifespan (3-8 months) (Terzibasi *et al*., 2008; Cellerino *et al*., 2016). Its rapid life-history strategy reflects adaptation to ephemeral savannah ponds (Reichard & Polačik, 2019). The species combines fast growth, early sexual maturation, high fecundity, and external fertilization, facilitating controlled breeding designs and experimental tests of parental effects (Cellerino *et al*., 2016). Despite its compressed lifespan, *N. furzeri* exhibits conserved cellular and molecular hallmarks of vertebrate ageing (Valenzano *et al*., 2015; Cellerino *et al*., 2016; Reuter *et al*., 2018). Age-dependent decline in global DNA methylation has already been reported in *N. furzeri* (Zupkovitz *et al*., 2021; Steiger *et al*., 2026). Turquoise killifish epigenetic clocks based on changes in DNA methylation perform well in age determination and expected lifespan prediction (Giannuzzi *et al*., 2024). In addition to that, the shift in DNA methylation in fins during ageing of the turquoise killifish corresponds relatively well with changes in ovaries and testes (Vrtílek *et al*., 2025).

Given the systematic changes in methylome with age across vertebrates and the evidence for incomplete DNA methylation marks erasure during embryogenesis in some fish, our aims were to test whether turquoise killifish: 1) DNA methylation changes systematically in ageing parents; 2) parental age affects offspring DNA methylation; and 3) the age-related DNA methylation changes in parents are transmitted to their offspring.

## Methods

We tested for inter-generational transfer of age-related epigenetic changes in a longitudinal experiment using a short-lived annual fish, the turquoise killifish. We analysed DNA methylation in parents at three pre-specified age-points as well as in their corresponding offspring (Figure 1). We retrieved information on methylation status across loci using reduced-representation sequencing approach (ddRAD) and examined both the overall proportion of DNA methylation and the overlap of methylated loci during ageing and between parents and their offspring. The experiment was replicated in three temporally separated blocks to increase the robustness of the results (for information on sample size per block see Table S1).

**Figure 1.**
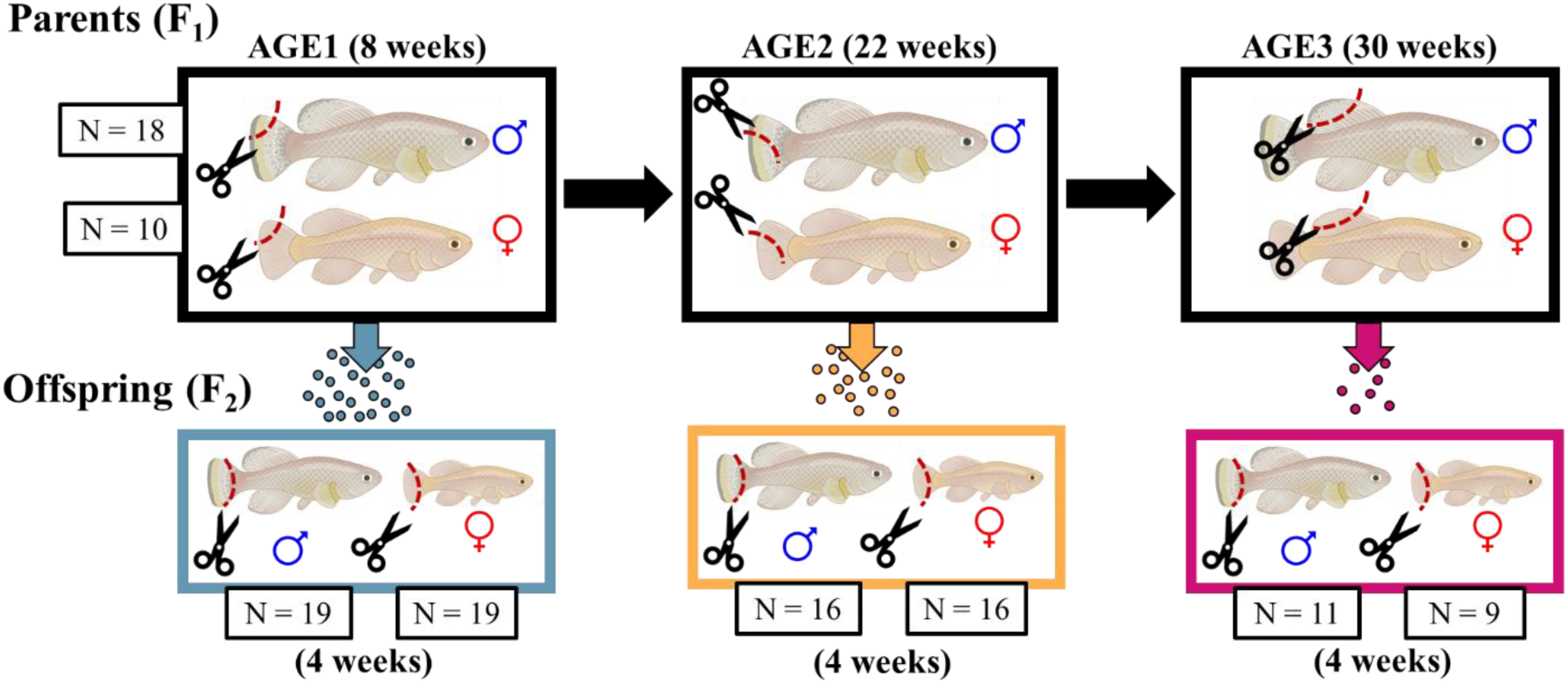
Sampling design for the analysis of DNA methylation between parents and their offspring in turquoise killifish. We collected non-regenerated fin tissue repeatedly from 10 dams and 18 sires at three pre-specified age-points. At the same time, we collected their clutches and reared offspring to test the inter-generational transmission of DNA methylation patterns. All offspring was sampled at the same age - 4 weeks post-hatching. Created using BioRender.

### Parental fish

For the experiment, we used wild-derived strain MZCS222 of the turquoise killifish (*Nothobranchius furzeri*) (Cellerino *et al*., 2016). We obtained the parental generation (F_1_) stock from mating of 34 different pairs of F_0_ generation. The clutches for F_1_ were incubated for 6-18 months, hatched and reared according to the published husbandry protocol (Polačik *et al*., 2016). Diet of the hatched fish initially consisted of *Artemia* sp. *nauplii* until 14 days post-hatching (dph) when weaned on adult diet of frozen bloodworms fed twice a day. At 10 dph, the parental fish were housed individually in common-garden conditions of a recirculation system (ZebTec, Tecniplast) with 27 °C (±1 °C), 1 mS/cm conductivity (reverse osmosis water mixed with NaCl) and 14-to-10 h light-to-dark regime. At the age of 4 weeks, when the sex of the fish was apparent, we assigned them into permanent families with three females (dams) and three males (sires) that were repeatedly mated with each other. There were in total 8 families in BLOCK1 and 6 in BLOCK 2 and BLOCK3 (see Vrtílek et al. 2026JEvB) but not all individuals were used in the present study.

### Fin sampling in parents

To capture potential changes in DNA methylation during ageing of parents, we collected fin samples repeatedly from 10 parental females (dams) and 18 males (sires) at three time-points. The AGE1 at 8 weeks corresponds to early adulthood (and 100% survivorship), AGE2 at 22 weeks to mid-age (with expected ca. 50% female mortality) and AGE3 at 30 weeks when marks of senescence are apparent (ca. 70% expected female mortality in the studied strain based survival data from (Žák & Reichard, 2021)). We timed the sampling according to females as the sex with lower life expectancy (Žák & Reichard, 2021).

The sampled individuals survived the whole experiment and also produced offspring during at least two AGE-samplings. At each AGE, we anaesthetised the fish with clove oil (Eugenol, 1 drop/0.1 L), cut a sample of their fin and stored it in −80 °C until DNA isolation. In parents, it was upper part of caudal fin at AGE1, lower part of the caudal fin at AGE2, and at AGE3, we cut the dorsal fin (Figure 1). We targeted non-regenerated tissue to avoid potential bias in age-related in DNA methylation changes due to regeneration (Hirose *et al*., 2013).

### Spawning and rearing offspring

We spawned the parental fish regularly twice a week and collected clutches for the offspring (F_2_) generation at the time of parental fin sampling (AGE1, AGE2, AGE3; see below, Figure 1). The fish were spawned in designated pairs outside their home tank in a 2-L plastic container with false mesh bottom (2×2 mm) for 1-1.5 h. Fertilized eggs (with visible perivitelline space) from each clutch were treated with methylene blue (FICHEMA, to final concentration ∼0.002 g/L) for 24 h after spawning, then placed on plastic Petri dish with moist peat and sealed with parafilm. We incubated the clutches for 5 months in 18.5 °C and darkness to simulate natural life cycle of annual killifish. To prepare embryos for hatching, we sterilized the eggs in a bath of peracetic acid solution (3 rounds alternating 5 min in autoclaved water and 5 min in a 0.5 mL/L solution of 4% of peracetic acid and hydrogen peroxide, Persteril 5, FICHEMA). The cleaned clutches were put onto a Petri dish with filter paper, sealed with parafilm and stored in 27 °C for three weeks. The increased temperature facilitates exit from embryonic diapause and completion of development (Polačik *et al*., 2016).

The offspring was hatched by watering the clutches with 22 °C tap water (0.45 mS/cm) and reared under the same conditions as their parents. At 5 dph, the juvenile offspring were moved to individual tanks in the recirculation system with the same setup as in their parents (see above). The offspring were fed twice a day initially with *Artemia* sp. *nauplii* and from 14 dph with chopped frozen bloodworms. We sampled offspring for caudal fin tissue at 4 weeks, when they were euthanized with clove oil overdose (5 drops/0.1 L). At this age, males showed their distinct colouration and females had round bellies with ovulated eggs. The tissue from offspring was always sampled at the same age, so that only their parents’ age differed (Figure 1).

### DNA extraction and sequencing

We have three separate blocks of samples for the parents and their offspring. For BLOCK1, the DNA was extracted using GeneJet purification kit according to the manufacturer protocol. For BLOCK2 and BLOCK3, genomic DNA was extracted from fin tissue using the cetyltrimethylammonium bromide (CTAB) protocol (Kovařík *et al*., 2000). DNA integrity then was verified by horizontal agarose-gel electrophoresis, and concentrations were determined with a NanoDrop spectrophotometer (Thermo Fisher Scientific) and a Qubit fluorometer (Thermo Fisher Scientific).

Reduced-representation sequencing to analyse DNA methylation, such as EpiRAD (Schield *et al*., 2016), has been successfully applied in fishes to detect environmentally and genetically structured DNA methylation variation, supporting their suitability for population-level epigenetic analyses (Lallias *et al*., 2021). We followed the protocol described by Vrtílek *et al*. (2025), which combines the traditional ddRAD-seq (double-digest Restriction-site Associated DNA sequencing) approach of (Peterson *et al*., 2012) with the 2RAD approach of (Bayona-Vásquez *et al*., 2019). This method combines double-digest RAD sequencing with restriction enzymes that differ in their sensitivity to DNA methylation, using a methylation-insensitive rare cutter (EcoRI) together with either the methylation-insensitive common cutter MspI or its methylation-sensitive isoschizomer HpaII. Because HpaII does not cut methylated CpG sites, comparison of libraries generated from the same samples allows inference of CpG methylation states at shared restriction loci. EpiRAD provides a cost-effective and scalable framework for population-level methylation analysis in non-model and emerging model systems (Peterson *et al*., 2012; Schield *et al*., 2016; Dimond *et al*., 2017).

Briefly, each of the total 174 DNA samples was digested in parallel with EcoRI–MspI and EcoRI–HpaII enzyme pairs. This generated paired RAD (EcoRI–MspI) and EpiRAD (EcoRI–HpaII) libraries for each sample. After digestion, adapters were ligated and DNA fragments were purified using SpriSelect beads (Beckman Coulter Life Sciences) (1 × volume ratio). The fragments were amplified using dual-indexed primers, and libraries were checked via horizontal agarose-gel electrophoresis. All libraries were pooled in equimolar amounts and concentrated with SpriSelect beads (1.2 ×). The pooled library was size-selected in tight mode, with an average fragment size of 340 base pairs (bp) using Pippin Prep (Sage Science). The final library was sequenced on the Illumina NovaSeq X Plus PE150 platform (Novogene Co. Ltd.), generating 150 bp paired-end reads. We sequenced 348 libraries in total. The quality of the raw sequencing reads was assessed using FastQC ver. 0.11.9 (Andrews, 2010) and summarized with MultiQC ver. 1.8 (Ewels *et al*., 2016). Inline barcodes, residual adapters, restriction enzyme recognition sites, and low-quality bases were trimmed using Skewer ver. 0.2.2 (Jiang *et al*., 2014). After trimming, the average number of reads per sample was 3.76 million for RAD and 4.45 million for EpiRAD libraries. The assembly was performed using ipyrad ver. 0.9.92 (Eaton & Overcast, 2020) with the *N. furzeri* MZM-0403 reference genome (GCF_027789165.1). To maximize locus recovery, the RAD and EpiRAD libraries from each sample were first concatenated prior to assembly, minimizing the potential loss of loci with low sequencing depth in individual libraries. This assembly generated 47 498 loci (19 681-35 420 loci per library, with a mean of 26 915 loci). Read counts for the reference loci in each RAD and EpiRAD library were extracted using SAMtools ver. 1.14 (Danecek *et al*., 2021). The *coverage* command was then used to calculate per-locus coverage statistics for each individual.

### Working with the resulting read counts

We standardized all libraries using “counts per million” (CPM) standardization (Lallias *et al*., 2021) to a common size to obtain the relative read representation of each locus in a library. The number of reads per locus was divided by the total number of reads mapped to the reference loci (total library size) and then multiplied by one million. To improve inference on whether a locus was methylated or not, we filtered loci with low relative read representation in the RAD libraries (restriction enzyme not sensitive to methylation, MspI). The minimum threshold was set to 15 reads per locus across all three experimental blocks based on block-specific analysis of the threshold impact on the number of retained loci and the proportion of DNA methylation. We used a value that maximised the number of retained loci for analysis per block while showing that the proportion of DNA methylation approximately plateaued (Figure S1). The same threshold value was also applied in our previous study of a similar type of samples (Vrtílek *et al*., 2025).

Filtering the RAD library separately per block resulted in 11 018 well-represented loci. The working dataset consisted of 0/1 data based on the combination of the RAD and EpiRAD libraries per each sample. The RAD library (from the methylation-non-sensitive MspI restriction enzyme) was used as a template of the loci present and the EpiRAD library (obtained using the methylation-sensitive HpaII restriction enzyme) then served for differentiation between methylated (1) and non-methylated state (0). A locus was defined as methylated when no reads were detected in the EpiRAD library. Such a strict approach may lead to an underestimation of overall DNA methylation because, at the tissue level, DNA methylation of a locus represents a continuous variable. However, accommodating continuous character of DNA methylation in our study would require introduction of a subjective sample-specific cut-off, thereby increasing analytical ambiguity (Dimond *et al*., 2017; Vrtílek *et al*., 2025). We therefore focus on the loci that were certainly methylated despite probably ignoring a large amount of DNA methylation.

### Data analysis

All data analyses were performed using R software ver. 4.4.1 (R Core Team, 2024).

First, we analysed the differences in proportion of methylated loci during parental ageing and in their offspring. The response variable was proportion of methylated loci in each sample. This was calculated as a mean per each sample from the well-represented loci (the 11 018 loci dataset) based on the 0/1 data, while ignoring NAs (as well-represented loci in a specific block could still be missing in the other blocks). We then fit and interpret the most complex model that included all the focal variables (AGE, BLOCK, SEX) and interaction between age and sex using mixed-effects models from “glmmTMB” package ver. 1.1.10 (Brooks, Mollie *et al*., 2017) accommodating the 0-1 data distribution. For offspring, the model also contained proportion of methylated loci of both the dam and the sire at the corresponding AGE-sampling (e.g., for offspring coming from parent’s AGE2, the proportion of DNA methylation recorded in their parents at AGE2). We accounted for the repeated sampling of parents by including individual parental ID as a random factor. In offspring, the model included the random effect of dam and sire ID.

The second step was to assess similarity in the loci methylation status across samples. We recovered 11 018 well-represented loci, but only 728 loci showed differential methylation across the 174 samples (i.e., at least one sample showed different methylation status at that locus compared to the others). Then, based on the methylation status (either 0 or 1) of the 728 variable loci, we computed the distances between samples using “dist.gene” function from “ape” package ver. 5.8 (Paradis & Schliep, 2019) and performed PCA (classical multi-dimensional scaling, function “cmdscale”) to see if there is a clustering among samples from different groups with regard to parental age. We then used PERMANOVA (function “adonis2” from package “vegan” ver. 2.7-2 (Oksanen *et al*., 2025) to examine which factor explained the most variation in DNA methylation loci status among the samples.

Finally, we extracted loci that became newly methylated (changed their status from 0 to 1) in parents between respective AGE-samplings (from AGE1 to AGE2, or AGE2 to AGE3) and checked if they were also methylated in offspring (either in the AGE2 or the AGE3 cohort). For example, we looked at newly methylated loci in a specific dam at AGE2 (compared to AGE1) and then whether her offspring at AGE2 showed DNA methylation at those specific loci. We only focused on newly methylated loci because of the low rate of DNA methylation in our samples. Because of the high prevalence of non-methylated loci, it did not make sense to inspect transfer of parental de-methylation to the offspring.

## Results

### 1. Offspring DNA methylation proportion was not affected by parental ageing or their proportion of methylated loci

In parents, the proportion of methylated loci increased with age in both dams and sires (log-odds of AGE2: 0.397±0.131 (estimate ± standard error), z = 3.032, P = 0.002), but AGE3-sampling was not different from the first (AGE1) sampling (0.185±0.136, z = 1.358, P = 0.175) (Table 1A, Figure 2A). There was no effect of parental age (Table 1B, Figure 2B), or parental proportion of DNA methylation (Table 1B) on offspring proportion of methylated DNA.

**Figure 2.**
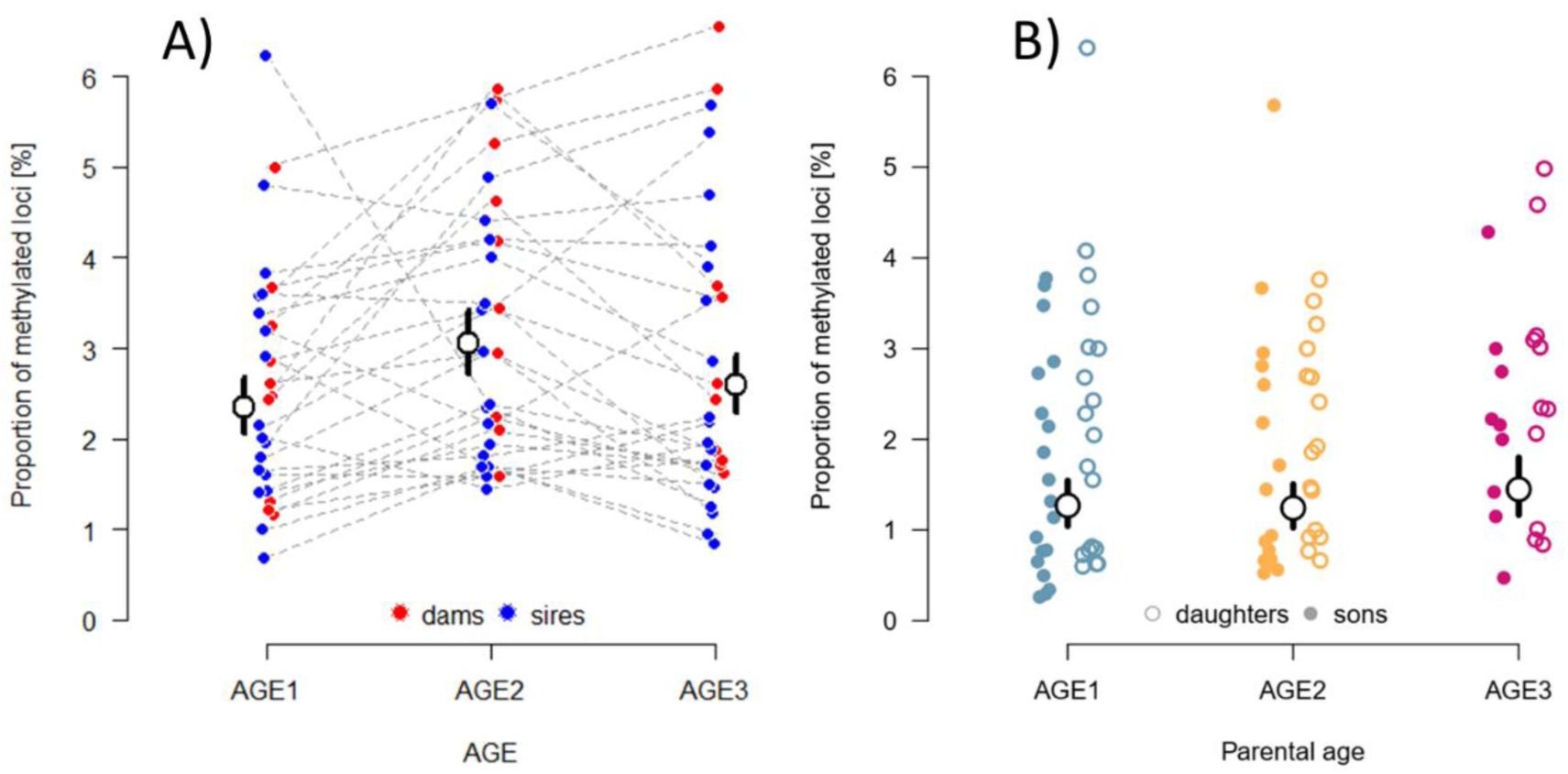
Proportion of methylated loci in parents (A) and their offspring (B). Parents (A) were sampled at 8 (AGE1), 22 (AGE2) and 30 weeks (AGE3), while the offspring were all sampled at 4 weeks after hatching. The large empty points show estimates for the AGE-groups (the effect of sex was not significant) along with 95% confidence intervals for the estimates (vertical lines). The smaller colour points show raw values that were jittered along the x-axis for better visibility. In the panel for parents (A), dashed grey lines illustrate repeated sampling of the same individuals.

**Table 1.** The analysis of the effect of parental age on the proportion of DNA methylation in parents (A) and their offspring (B). The terms highlighted **in bold** were statistically significant (at α = 0.05). S.E. – standard error of the estimate. The intercept represents dams from AGE1 sampling of BLOCK1 in A) and daughters from AGE1 sampling of BLOCK1 in B).

| A) | Term | Estimate | S.E. | z | P |
| --- | --- | --- | --- | --- | --- |
|  | *Fixed effects:* |  |  |  |  |
|  | Intercept | -3.315 | 0.108 | -30.822 | <0.001 |
|  | **AGE2** | **0.397** | **0.131** | **3.032** | **0.002** |
|  | AGE3 | 0.185 | 0.136 | 1.358 | 0.175 |
|  | SEX(sires) | 0.072 | 0.131 | 0.551 | 0.581 |
|  | **BLOCK2** | **-0.468** | **0.094** | **-4.964** | **<0.001** |
|  | **BLOCK3** | **-0.875** | **0.092** | **-9.508** | **<0.001** |
|  | AGE2: SEX(sires) | -0.255 | 0.168 | -1.520 | 0.128 |
|  | AGE3: SEX(sires) | -0.169 | 0.174 | -0.969 | 0.333 |
|  | *Random effect:* |  |  |  |  |
|  | ID | 0.007 |  |  |  |
| B) | Term | Estimate | S.E. | z | P |
|  | *Fixed effects:* |  |  |  |  |
|  | Intercept | -3.284 | 0.229 | -14.352 | <0.001 |
|  | AGE2 | -0.059 | 0.140 | -0.422 | 0.673 |
|  | AGE3 | 0.006 | 0.145 | 0.044 | 0.965 |
|  | SEX(sons) | -0.242 | 0.127 | -1.914 | 0.056 |
|  | dam DNA methylation | -0.015 | 0.041 | -0.368 | 0.713 |
|  | sire DNA methylation | -0.054 | 0.045 | -1.204 | 0.229 |
|  | **BLOCK2** | **-0.943** | **0.194** | **-4.852** | **<0.001** |
|  | **BLOCK3** | **-1.235** | **0.203** | **-6.078** | **<0.001** |
|  | AGE2: SEX(sons) | 0.074 | 0.189 | 0.391 | 0.696 |
|  | AGE3: SEX(sons) | 0.255 | 0.202 | 1.258 | 0.208 |
|  | *Random effects:* |  |  |  |  |
|  | dam ID | <0.001 |  |  |  |
|  | sire ID | 0.023 |  |  |  |

### 2. Offspring methylation pattern did not correspond with their parents’ age

The variation in methylation status across loci in offspring was not associated with their parents’ AGE-group (Figure 3). The variation explained by the age of parents or offspring sex was marginal (PERMANOVA: R^2^ = 0.019 and 0.009, for age and sex, respectively) compared to the effect of experimental block (R^2^ = 0.108).

**Figure 3.**
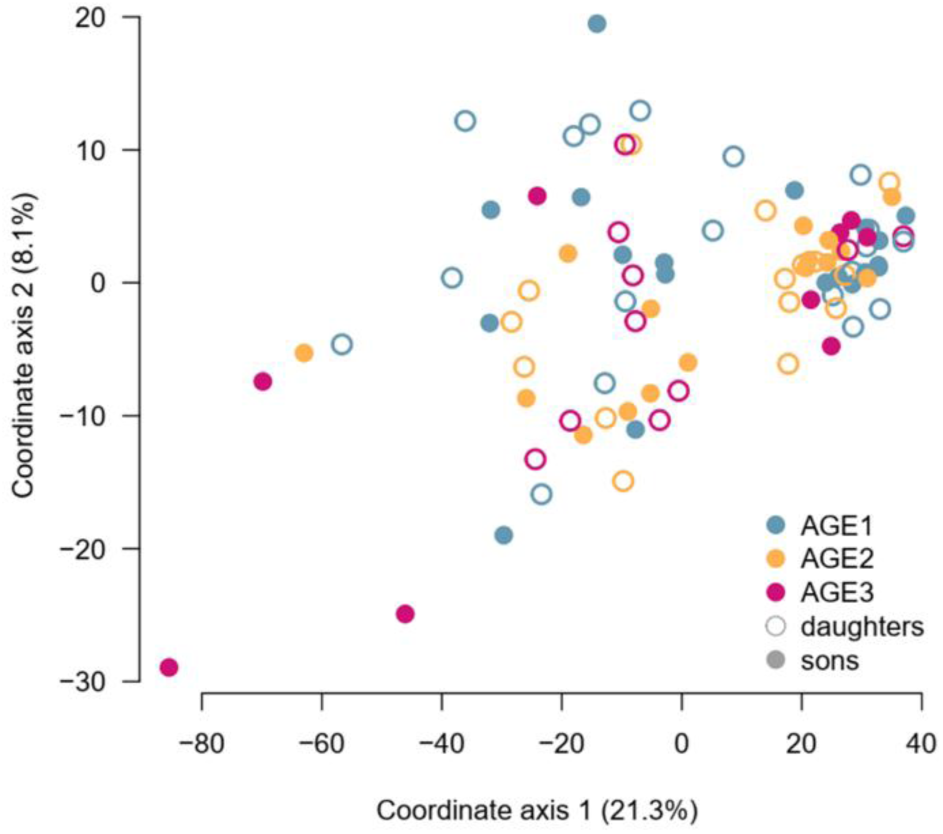
Similarity of DNA methylation among offspring based on the 728 differentially methylated loci. Offspring colour-code refers to the parental age in two-dimensional space based on PCA of DNA methylation status across the loci. Each point represents one offspring (full for male offspring and empty for female offspring). Axis 1 and axis 2 explain 21.3% and 8.1% of the variation, respectively. Two extreme values were excluded for better illustration of the overall pattern. The effect of different block can be seen in Figure S2.

### 3. Parental age-related changes in methylation did not transmit to offspring

We identified 728 differentially methylated loci across parents and their offspring (methylation status of a locus being different in at least one sample). These loci thus potentially contained information related to parental ageing or parent-offspring relationships.

In parents (dams and sires analysed separately), there were a lot of loci overall that changed their methylation status during ageing (in total 226-396 loci in individual dams and 358-395 in individual sires, out of the total 728 loci). This means that almost one third to more than half of the differentially methylated loci were involved in either new methylation or de-methylation between AGE1 and AGE2, or AGE2 and AGE3 samplings. However, the recorded age-related methylation changes were not widely shared across individuals. There was only a single locus where a *de novo* methylation occurred across more than half of dams between AGE1 and AGE2. No other change in locus methylation status was prevalent in more than 50% of dams or sires.

Given the low amount of shared methylation associated with ageing in parents, we decided to analyse the inter-generational transmission of methylation patterns individually, for each dam, or sire, and their offspring. The number of newly methylated loci in dams at AGE2 and AGE3 varied from 30 to 124 and from 19 to 108, respectively. In sires, it was 16-129 loci at AGE2 and 12-119 at AGE3 (Table S2). On average, only few of these loci were also methylated in at least half of their offspring from the corresponding AGE (5 in AGE2 of dams, 5 in AGE2 of sires, 7 in AGE3 of dams and 6 in AGE3 of sires) (for details see Table S2). In the end, there were only few individual loci affected at each parent-offspring combination and the identity of the focal loci varied across the combinations. This means we were not able to identify a common DNA methylation trace from ageing parents to their offspring at respective AGE-points.

## Discussion

We tested whether the changes in DNA methylation during ageing are transmitted to offspring in a model short-lived fish, the turquoise killifish. Longitudinal sampling of the parental fish revealed an increase in the proportion of methylated loci that was only apparent in the mid-age. More importantly, DNA methylation profile of their offspring did not correspond with the ageing dynamics nor with the individual patterns recorded in parents. The absence of parental imprint through DNA methylation in the turquoise killifish is probably due to early (embryonic) epigenetic erasure (Fellous *et al*., 2018; Wang & Bhandari, 2019).

### Changes in DNA methylation during parental ageing

Overall DNA methylation increased in parents at mid-age (AGE2 sampling) but then, when becoming older (at AGE3), the proportion of methylated loci was similar to their young phase (AGE1). In a similar but cross-sectional experiment with the turquoise killifish, we recorded an increase in the proportion of methylated loci towards the old age (AGE3) (Vrtílek *et al*., 2025). The transient increase in methylation at AGE2 and decline at AGE3 may reflect a period of epigenetic remodelling during mid-life, followed by partial restoration or further reorganisation at older age. Similar non-linear patterns, including a return towards a younger methylation profile later in life, have been reported in mice and in the turquoise killifish (Olecka *et al*., 2024; Steiger *et al*., 2026). Collecting more time-points in the future could explain what is really occurring at the global methylation level during turquoise killifish ageing and help to resolve the temporal locus-specific changes underlying this pattern. Why the overall DNA methylation increased first (AGE1-AGE2) and then declined again (AGE2-AGE3) is an open question.

Global methylation represents the net outcome of many locus-specific changes. Age-related shifts in DNA methylation result from both stochastic epigenetic drift and active de-/re-methylation (Le Clercq 2023). Local changes may occur in the opposite direction to the global pattern, thereby obscuring the overall methylation pattern (Sen *et al*., 2016). Using the repeated sampling of the non-regenerated fins, we demonstrated substantial variation among individual trends in the overall proportion of methylated loci during ageing (Figure 2A). Certain part of the variation among individuals observed here can be ascribed to the stochastic epigenetic drift in aging (Hernando-Herraez *et al*., 2019; Nick Weber *et al*., 2024).

After filtering and data processing, 728 differentially methylated loci were retained for locus-specific analysis of similarity in DNA methylation. Age-related changes at these loci showed little overlap among individuals, indicating that most detectable changes within this reduced representation screen of the genome were not consistently shared across the population. We therefore did not identify a common set of age-associated loci in the present dataset. Importantly, this should not be interpreted as evidence that turquoise killifish lack reproducible epigenetic ageing signatures. Such signatures and accurate epigenetic clocks have been demonstrated using genome-wide datasets (Giannuzzi et al., 2024), and more generally, DNA methylation dynamics provide a reliable basis for age estimation (Le Clercq et al., 2023). However, robust epigenetic clock development requires dense genomic coverage and sampling across a continuous age range (Mayne *et al*., 2020; Giannuzzi *et al*., 2024). Our study was not designed for this purpose; instead, it provides longitudinal evidence that global and locus-specific methylation trajectories can vary markedly among individuals and may change non-linearly over the adult lifespan.

### Parental effects on DNA methylation in offspring

The age-related variation in DNA methylation in parents did not transmit to their offspring. We were not able to find a common ageing methylation pattern in parents (between AGE1 and AGE2 and between AGE2 and AGE3) that would be shared with their offspring (either in AGE2 or AGE3 clutches). Our results therefore suggest that age-associated DNA methylation changes are largely reset between generations, most likely through methylation erasure during embryogenesis. This is consistent with the widespread methylation erasure reported during embryogenesis in many vertebrates (Skvortsova *et al*., 2018; Wang & Bhandari, 2019; Fitz-James & Cavalli, 2022). It is also supported by developmental expression patterns of dnmt and tet genes in the turquoise killifish, which suggested that molecular machinery required for embryonic DNA methylation remodelling is active during early development (Zupkovitz *et al*., 2021). In contrast, studies on zebrafish indicate that embryonic DNA methylation reprogramming is incomplete and that offspring methylation patterns reflect paternal methylome (Potok *et al*., 2013; Skvortsova *et al*., 2019).

It is true that we only sampled offspring when they were adult, thus we do not possess direct evidence for the methylation erasure during embryonic development. However, two waves of demethylation have been reported in medaka (during primordial germ and at cell stage 0-25dpf) (Wang & Bhandari, 2019, 2020). At least one wave has been reported during embryogenesis in mangrove rivulus (Fellous *et al*., 2018), a closely related species to the turquoise killifish. These findings raise question to what extent can zebrafish results be generalized to other fish species. Future work should therefore directly investigate DNA methylation dynamics during embryogenesis in turquoise killifish and other non-model fish species to test how widely prevalent the DNA methylation erasure actually is and what are the developmental consequences.

From an evolutionary perspective, our results support the idea that extensive embryonic methylation reprogramming may protect offspring from inheriting age-associated epigenetic deterioration. Obviously, accumulation of adverse aspects of senescence over generations should be somehow selected against.

### Limitations of our study

We captured a relatively low proportion of methylated loci (maximum 6.5% of focal loci were methylated) compared to similar studies. In fish, including turquoise killifish (Steiger *et al*., 2026), the global levels of CpG methylation are typically 60-80% (Fellous *et al*., 2018; Skvortsova *et al*., 2018; Bélik & Silvestre, 2026; He *et al*., 2026). First, we used reduced representation sequencing focused on the CpG dinucleotides. This resulted in non-random and low coverage sampling of the genome. Second, and probably the main reason for the low methylation rate recorded, was the way that we processed the read-count data into the presence/absence of locus methylation.

The EpiRAD sequencing does not identify the methylation status of retrieved loci directly as through bisulfite sequencing, for example (Giannuzzi *et al*., 2024; Bélik & Silvestre, 2026). For a genetically heterogeneous population, a second – template - library is necessary for identification present loci along with the methylation-sensitive library. The number of reads in the template library then also allows estimation which locus was methylated (Dimond *et al*., 2017). The use of EpiRAD thus requires two fundamental data processing steps – loci filtering (in the template library) and methylation estimation itself (methylation-sensitive library).

With the first step, we focused on loci that were represented with a given sequencing minimum depth across all samples to maximise the generalization/objectivity of our inference (Mayne *et al*., 2020; Liebl *et al*., 2021; Tangili *et al*., 2025). The minimum sequencing depth (number of reads per locus in the template library) was used to provide sufficient contrast between unmethylated and methylated state by removing loci with very low read numbers from the template library. This allowed us to streamline and standardize our working pipeline.

The second important step in our pipeline was the threshold criterion to decide locus methylation status. The criterion for a locus to be considered methylated was zero reads in the methylation-sensitive library. This strict threshold inevitably led to ignoring some loci that were also largely, but not completely, methylated – these were labelled as non-methylated. The methylation status of a certain locus in a tissue is represented by a continuum (proportion) not by 0/1 methylation status. While it is possible to approach the dataset more inclusively by setting the limit to a ratio between the number of reads in the template and methylation-sensitive library (Dimond *et al*., 2017), it would introduce non-systematic sample-specific bias in the results as described by (Vrtílek *et al*., 2025). Strict threshold for identification of methylated loci was therefore a straightforward and more defendable option for retrieving robust information on methylation in at least part of the methylated loci.

## Conclusion

We tested for modulation of epigenetic inheritance due to parental ageing in a short-lived fish, the turquoise killifish. The recorded changes in parental DNA methylation were non-linear, individual-specific, and not reflected in the offspring. While there is robust research available on epigenetic ageing in the turquoise killifish (Giannuzzi *et al*., 2024), the transmission of these changes to the next generation with potential accumulation of negative effects of senescence is still an open question. More analysis is warranted especially into the taxonomic extent of the methylation erasure during early embryogenesis as this aspect was beyond the scope of the present study.

## Author contribution

**Malahat Dianat** – Investigation, Data curation, Formal analysis, Writing – review and editing

**Lisandrina Mari** – Conceptualization, Investigation, Writing – review and editing

**Dagmar Čížková** – Conceptualization, Methodology, Supervision, Writing – review and editing

**Milan Vrtílek** – Conceptualization, Funding acquisition, Investigation, Data curation, Formal analysis, Visualization, Writing – original draft, Writing – review and editing

## Acknowledgements

We would like to acknowledge the Mendel Centre for Plant Genomics and Proteomics at CEITEC, Masaryk University for the DNA isolation of BLOCK2 and BLOCK3 samples. We appreciate the help from Mahdi Mahmoudi (ISTA, Austria) during the data analysis. Computational resources were provided by the e-INFRA CZ project (ID:90254), supported by the Ministry of Education, Youth and Sports of the Czech Republic. This study was supported by the Czech Science Foundation (22-21198S).

## Supplementary information

**Table S1.** The distribution of samples between the three experimental blocks. The counts show for each AGE-sampling the number of collected samples per block separated by a slash. Total gives total number of individuals sampled (parents were sampled repeatedly).

| Parents | AGE1 | AGE2 | AGE3 | Total |
| --- | --- | --- | --- | --- |
| dams | 5/2/3 | 5/2/3 | 5/2/3 | 10 |
| sires | 6/5/7 | 6/5/7 | 6/5/7 | 18 |
| Offspring | AGE1 | AGE2 | AGE3 | Total |
| daughters | 11/3/5 | 10/2/4 | 9/0/2 | 46 |
| sons | 10/4/5 | 9/2/5 | 6/0/3 | 44 |

**Table S2.**
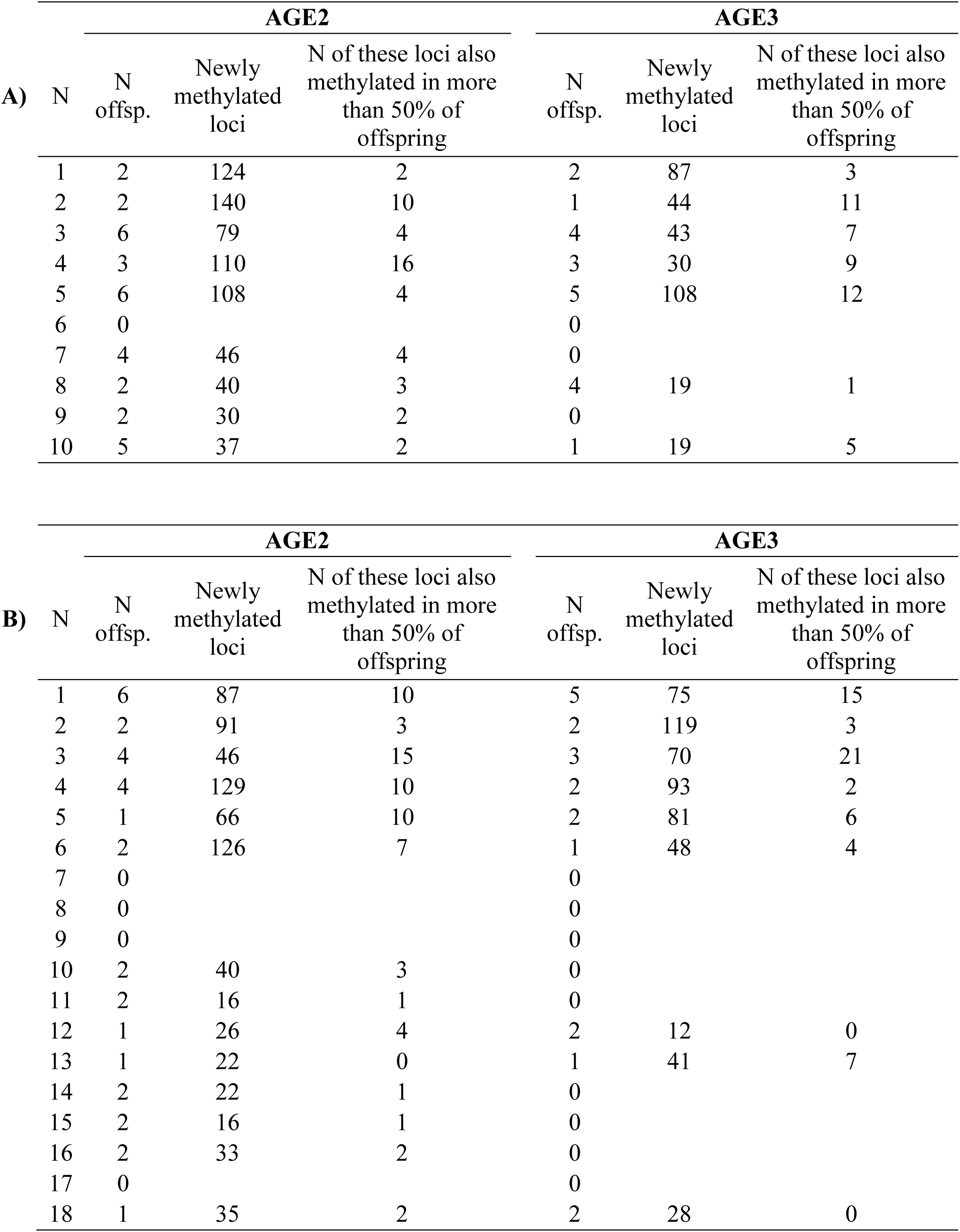
The overview of locus-specific analysis of DNA methylation transmission from parents (A - dams, B - sires) onto their offspring. The analysis was performed for dams and sires separately and split by AGE – AGE2 for AGE1-AGE2 transition and AGE3 for AGE2-AGE3 transition. The total number of differentially methylated loci was 728.

**Figure S1.**
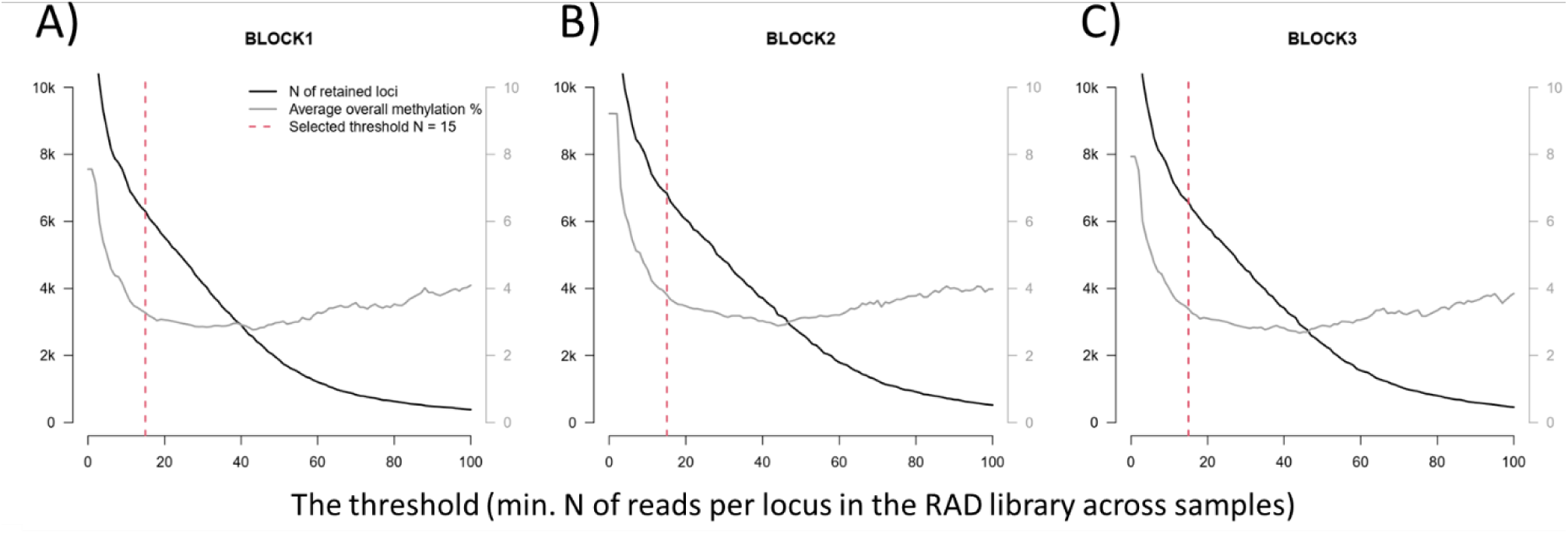
The impact of selected filtering threshold on the number of retained loci and average proportion of methylated loci across samples in the respective experimental blocks.

**Figure S2.**
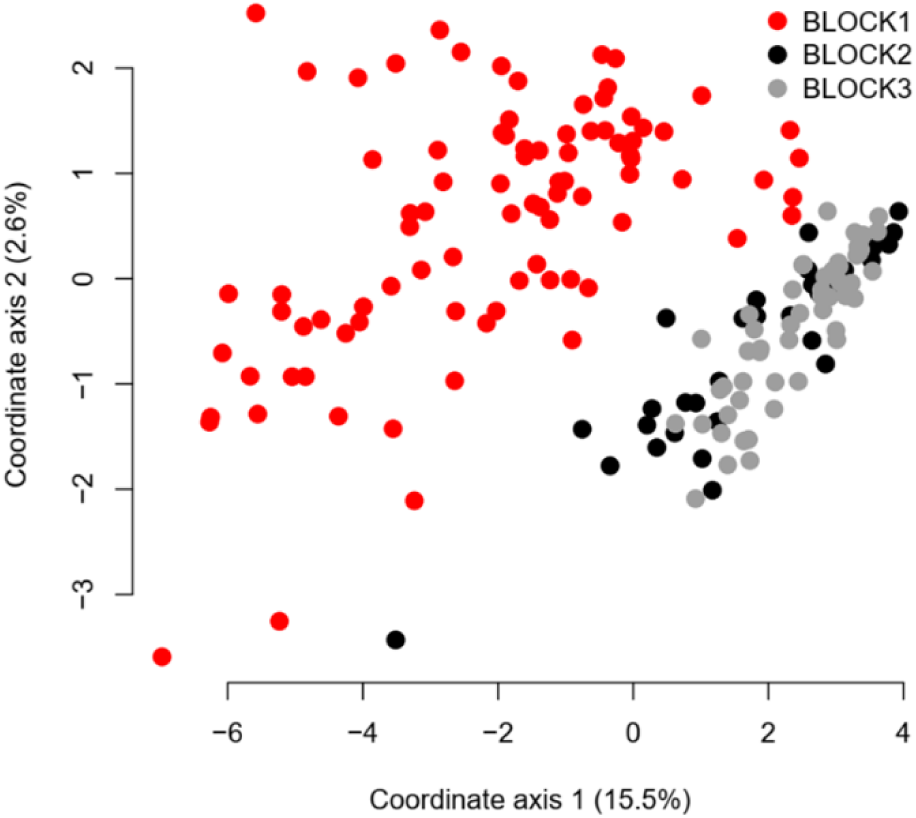
Similarity analysis of DNA methylation across all samples (parents and offspring together) colour-coded according the experimental blocks.

